# Pharmacological stress exposes hidden allelic background effects in genetic interaction screen normalisation

**DOI:** 10.64898/2026.05.21.726896

**Authors:** Rowshan Islam, Olga Xintarakou, Charalampos Rallis

## Abstract

Synthetic Genetic Array (SGA) analysis comprises the high-throughput crossing of a query deletion strain against a genome-wide deletion library to score fitness interactions in thousands of double mutants. SGAs have produced comprehensive genetic interaction maps in yeasts and have emerged as a leading platform for pharmacogenomics: mapping genetic modifiers of drug response, identifying synthetic lethal targets and illuminating mechanisms of drug action and resistance. We have previously demonstrated that in fission yeast, the *ade6* mutant is functionally neutral relative to the parental library and can serve as a standard negative control for SGA screens. Here, while we confirm our previous observation, we show that this neutrality fails under pharmacological stress. Using Torin1, an ATP-competitive TOR kinase inhibitor, we demonstrate that the *ade6* SGA fitness profile diverges from that of the parental library in a dose-dependent and genomically widespread manner. At 2 μM Torin1 only 12.2% of scored genes exceed a 1.5-fold fitness difference between backgrounds; at 3 μM this proportion rises to 43.2% -a 3.5-fold increase driven by qualitative reorganisation of the genetic interaction landscape rather than simple scaling of pre-existing differences. Gene ontology analysis of divergent genes implicates autophagy, iron starvation responses, central carbon metabolism, and vesicle trafficking, consistent with TOR-regulated nutrient adaptation being differentially affected by the *ade6-M210/M216* point mutations in the library versus the *ade6* null in the SGA control. Our results have implications in fission yeast and beyond and we propose solutions towards reliable retrieval of genetic interactions.

## Introduction

Synthetic Genetic Array (SGA) analysis maps the fitness consequences of pairwise gene deletions at genome scale, making it one of the most powerful tools in functional genomics ^1,2^. Originally developed for *Saccharomyces cerevisiae* (budding yeast)^3^, SGA automates the generation of double mutants to identify functional relationships between genes, such as synthetic lethality and suppression. Extending this technology to *Schizosaccharomyces pombe* (fission yeast)^4,5^, enabled the discovery of conserved eukaryotic pathways^4^ and the genetic underpinnings of fundamental cellular processes^2,6–8^. The technique has been extended to trigenic interaction studies^9–11^ in both budding and fission yeasts and in combination with drugs^12,13^.

SGA involves sequential steps of mating, sporulation, and haploid selection to generate combinatorial double mutants, which are then screened for growth phenotypes to map genetic interactions. The power of this approach for large-scale interaction mapping has been most strikingly illustrated by construction of comprehensive genetic interaction networks in *S. cerevisiae*^1^. These networks not only reveal functional relationships but also systematically annotate gene function, identify pathway components, and illuminate the architecture of complex traits^2,14^. In *S. pombe*, the adaptation of SGA is especially powerful because fission yeast shares fundamental eukaryotic features with humans, including chromosomal architecture, cell cycle regulation, and RNA-processing machinery, making it an ideal model for identifying conserved genetic interactions and translating network-level insights to higher organisms^15^. The SGA approaches in yeasts have inspired expanding genome-scale genetic interaction networks for human cell lines^16^.

The integration of SGA with drugs and compounds provides a powerful route to identifying genetic determinants of drug response. Synthetic lethality screens have been instrumental in discovering drug targets and resistance mechanisms, particularly in cancer^17^. In yeast models, chemical-genetic SGA screens probe interactions between compounds and gene deletions, enabling prediction of drug synergies and informing combination therapies. Critically, however, the validity of pharmacogenomic SGA depends on the suitability of the chosen normalisation control under the specific drug condition used.

The *ade6* SGA is performed by mating the haploid deletion library with an *ade6Δ* query strain and has previously been used as a negative control for SGA screens^9,13,18^. The Bioneer haploid deletion library carries the *ade6-M210* or *ade6-M216* point-mutation alleles^19^, which disrupt *ade6* function. Introducing an *ade6Δ* through SGA adds a second deletion event without introducing a new adenine-biosynthesis defect in adenine-supplemented medium, making it a practical negative control. Critically, however, the *ade6-M210/M216* alleles are point mutations that retain the *ade6* locus and may differ from a complete null in their metabolic and regulatory consequences. This latent difference is inconsequential under standard growth conditions but, as we show here, is exposed by TOR inhibition in a dose-dependent manner, calling into question the use of the *ade6* SGA as a normalisation control in pharmacogenomic screens and proposition of alternative methods going ahead.

## Results

We compared two genetic backgrounds -the haploid single-deletion library and the *ade6* SGA background (generated by crossing the library with an *ade6*Δ query strain) -each profiled under three conditions: (i) no drug, (ii) 2 μM Torin1 (T2), and (iii) 3 μM Torin1 (T3), yielding nine pairwise contrast signatures (Figure 1A). We then: (i) examined the global structure of these contrasts by Pearson correlation, hierarchical clustering, and PCA; (ii) assessed the direct baseline difference between backgrounds without drug; (iii) compared within-background Torin1 responses; (iv) quantified between-background divergence under Torin1; and (v) applied a difference-in-differences framework to isolate the interaction between drug effect and genetic background.

**Figure 1.**
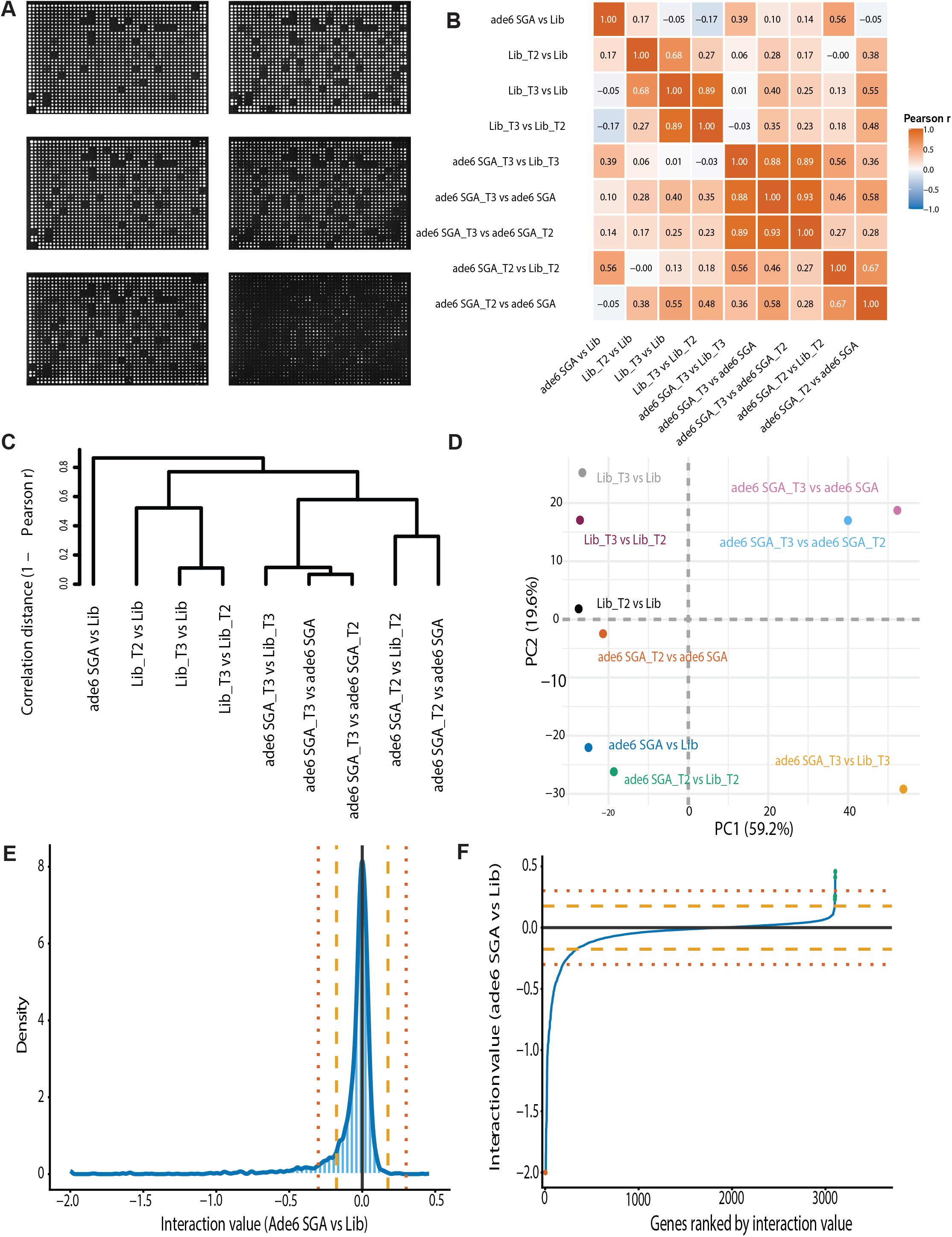
Global overview of contrast structure and baseline comparison of *ade6* SGA with the library. (A) Left and right panels show the corresponding plates of the haploid deletion library and *ade6* SGA background; from top to bottom, untreated, 2 μM Torin1-treated, and 3 μM Torin1-treated plates. (B) Pearson correlation heatmap showing the relationships among the nine contrast signatures. The numerical values in each cell indicate Pearson correlation coefficients. (C) Hierarchical clustering dendrogram generated using correlation distance (1−Pearson r). (D) Principal component analysis of the same contrast signatures. Percentages on the axes indicate the proportion of variance explained by each principal component. (E) Distribution of interaction values for *ade6* SGA vs Lib across all analysed genes. The solid black vertical line marks 0, indicating no difference between the *ade6* SGA background and the library. Dashed orange lines indicate interaction values of ±0.176 [±log_10_(1.5)], and dotted orange lines indicate interaction values of ±0.301 [±log_10_(2)]. (F) Ranked interaction values for *ade6* SGA vs Lib. Genes are ordered from the most negative to the most positive interaction value. The solid black horizontal line marks 0, dashed orange lines indicate the ±1.5-fold threshold, and dotted orange lines indicate the ±2-fold threshold.

### Global structure of the contrasts

To evaluate whether Torin1 treatment reshapes the relationship between the two genetic backgrounds, we applied Pearson correlation, hierarchical clustering, and Principal Component Analysis (PCA) to all pairwise contrast signatures (Figure 1B-D). The correlation heatmap (Figure 1B) showed that library-based Torin1 contrasts cluster tightly (strongest: Lib_T3 vs Lib with Lib_T3 vs Lib_T2, r = 0.89), as do the *ade6*-associated T3 contrasts (*ade6* SGA_T3 vs *ade6* SGA with *ade6* SGA_T3 vs *ade6* SGA_T2 showing the highest correlation in the dataset, r = 0.93). Critically, the high-dose between-background contrast (*ade6* SGA_T3 vs Lib_T3) aligned strongly with the *ade6*-associated T3 contrasts (r = 0.88–0.89) yet was not correlated with the corresponding library-only contrasts (Lib_T3 vs Lib: r = 0.01; Lib_T3 vs Lib_T2: r = −0.03). This dissociation establishes that, at high Torin1 doses, the *ade6* SGA background diverges from the parental library in a manner driven primarily by the *ade6* background itself, not by the drug. Hierarchical clustering (Figure 1C) corroborated this structure: library-centred contrasts co-clustered tightly, while the *ade6*-associated T3 contrasts and the between-background comparison formed a distinct module separated from both the library-centred and untreated contrasts.

PCA (Figure 1D) provided a consistent picture. PC1 (59.2% of variance) separated the *ade6*-associated T3 contrasts from all library-centred and lower-dose contrasts, with the dominant source of variation being the emergence of a high-dose *ade6*-associated Torin1 response. PC2 (19.6%) further resolved between-background from within-background comparisons. Notably, *ade6* SGA_T3 vs Lib_T3 separated markedly from *ade6* SGA_T2 vs Lib_T2 along PC1, confirming that 3 μM Torin1 does not simply scale the 2 μM between-background difference but qualitatively reshapes the *ade6*-associated interaction landscape.

### Baseline comparison between haploid deletion library and *ade6* SGA

Baseline comparison between the *ade6* SGA control and the haploid deletion library in the absence of drug was performed to establish genome-wide equivalence before any pharmacological perturbation was applied. In SGA terminology, an interaction value near zero indicates that the fitness of the double mutant (*ade6*Δ; *gene*Δ) is indistinguishable from that expected from the single mutant alone; a distribution sharply centred on zero across the whole genome therefore means the *ade6* deletion contributes no second-mutation effect. Consistent with this expectation, interaction values for *ade6* SGA vs Lib were sharply centred near zero (Figure 1E–F), confirming that the two backgrounds are broadly equivalent under standard growth conditions and that the *ade6* SGA is a suitable negative control for conventional screens. Nonetheless, the distribution was not perfectly symmetric: the negative tail was broader and more extreme than the positive tail, and the ranked interaction plot (Figure 1F) revealed a subset of genes with pronounced negative deviations alongside fewer positive outliers. This residual asymmetry, limited under basal conditions but already pointing to a non-neutral allelic background at a minority of loci, provides essential context for interpreting the far greater divergence that emerges under TOR inhibition.

### Within-background and between-background drug-response comparison

To compare Torin1 responses between the haploid deletion library and the *ade6* SGA background, we concentrated on gene-level fitness changes in each background at each dose (Figure 2A, 2B). If the two backgrounds respond equivalently to the drug, genes should cluster tightly along the identity line (y = x) with few deviating by more than the ±log_10_(1.5) threshold used here to define meaningful fitness divergence, corresponding to a 50% fitness difference between backgrounds. At 2 μM Torin1, the two backgrounds showed moderate genome-wide correspondence (r = 0.56, RMSE = 0.163, regression slope 1.13), with most genes falling reasonably close to the identity line; 357 of 2,938 scored genes (12.2%) exceeded the divergence threshold. At 3 μM Torin1, this picture changed markedly: correspondence fell substantially (r = 0.39, RMSE = 0.269, regression slope 0.69), the scatter widened and 1,131 of 2,616 scored genes (43.2%) exceeded the threshold (a 3.5-fold increase in divergent genes relative to the lower dose). The change in regression slope from 1.13 to 0.69 indicates that the nature of the divergence also shifted qualitatively: rather than a proportional amplification of a consistent quantitative difference, increasing TOR inhibition produces a fundamentally different pattern of relative fitness effects across the genome. These results demonstrate that at pharmacologically active Torin1 concentrations the *ade6* SGA background cannot be treated as a neutral proxy for the parental library, and that this failure worsens non-linearly with dose.

**Figure 2.**
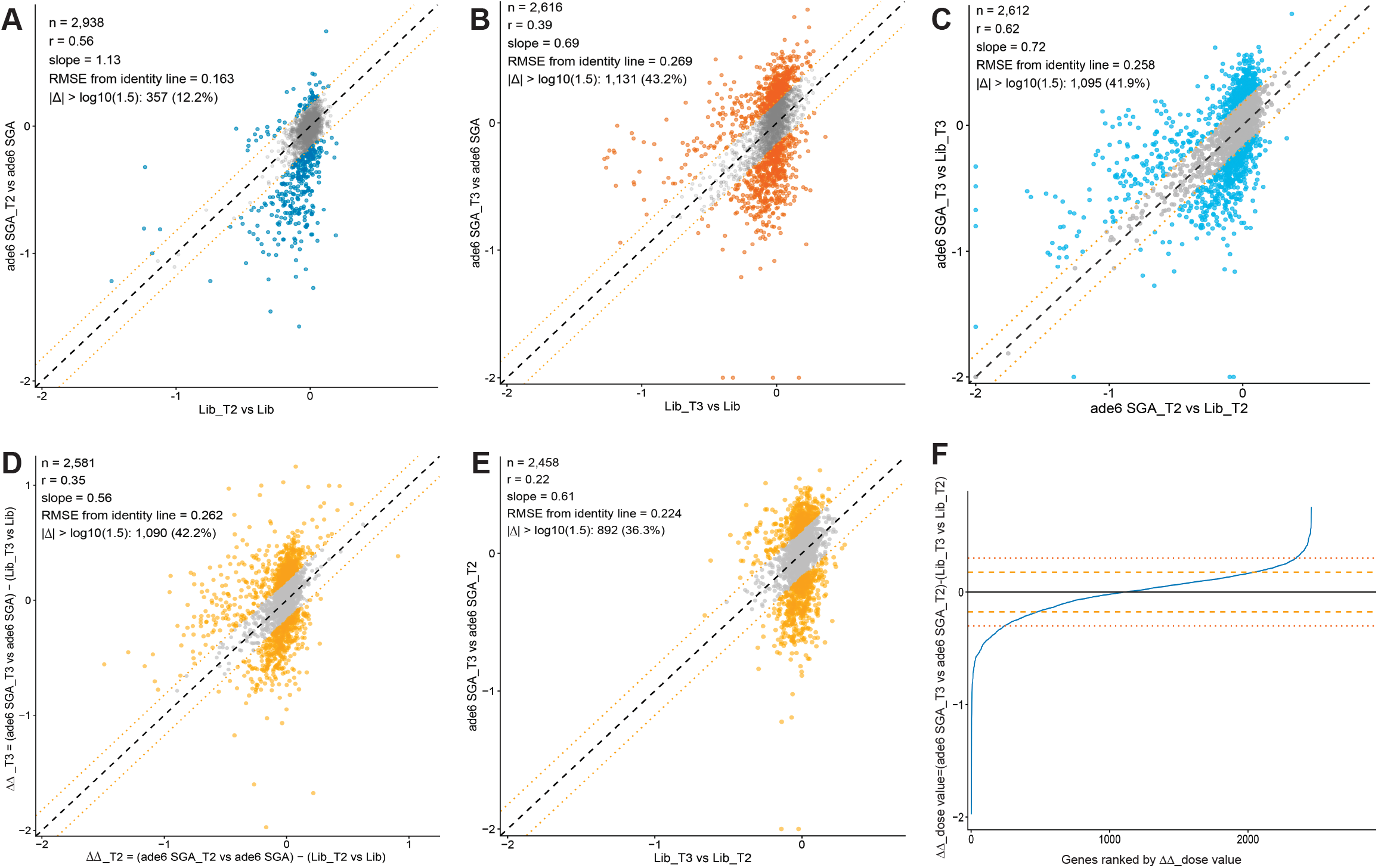
Comparison of Torin1 responses in the haploid deletion library and *ade6* SGA. (A) Identity scatter plot comparing the Torin1 response in the haploid deletion library and *ade6* SGA background at 2 μM Torin1. (B) Identity scatter plot comparing the Torin1 response in the haploid deletion library and *ade6* SGA background at 3 μM Torin1. (C) Identity scatter plot comparing the between-background contrasts *ade6* SGA_T2 vs Lib_T2 and *ade6* SGA_T3 vs Lib_T3. (D) Identity scatter plot comparing ΔΔ_T2 and ΔΔ_T3, where ΔΔ_T2 = (*ade6* SGA_T2 vs *ade6* SGA) - (Lib_T2 vs Lib) and ΔΔ_T3 = (*ade6* SGA_T3 vs *ade6* SGA) − (Lib_T3 vs Lib). (E) Identity scatter plot comparing the dose-step responses Lib_T3 vs Lib_T2 and *ade6* SGA_T3 vs *ade6* SGA_T2. (F) Ranked ΔΔ_dose values, where ΔΔ_dose = (*ade6* SGA_T3 vs *ade6* SGA_T2) − (Lib_T3 vs Lib_T2). Genes are ordered from the most negative to the most positive value. For panels A-E, each point represents a gene. The dashed black line indicates the identity relationship (y = x), and the dotted orange lines indicate a deviation of ±log_10_(1.5) from the identity line. For panel F, the solid black horizontal line marks 0, dashed orange lines indicate ±log_10_(1.5), and dotted orange lines indicate ±log_10_(2).

Comparing the between-background contrasts directly (*ade6* SGA_T2 vs Lib_T2 against *ade6* SGA_T3 vs Lib_T3; Figure 2C), the two doses showed moderate correlation (r = 0.62, RMSE = 0.258) with a slope of 0.72, indicating that between-background divergence does not scale proportionally with dose. Of 2,612 genes, 1,095 (41.9%) exceeded the ±log_10_(1.5) threshold, confirming that the *ade6* SGA background diverges from the parental library in a dose-dependent and non-proportional manner under Torin1 treatment.

### Difference-in-differences and dose dependence response analysis to Torin1

Direct between-background comparisons (*ade6* SGA_Tx vs Lib_Tx) reflect both pre-existing background differences and drug-induced divergence without separating the two. To disentangle these, we applied a difference-in-differences (DiD) framework, which subtracts the within-background Torin1 response of the library from the within-background Torin1 response of the *ade6* SGA background, yielding a metric that specifically captures the background-specific component of the drug effect: ΔΔ_T2 = (*ade6* SGA_T2 vs *ade6* SGA) − (Lib_T2 vs Lib) and ΔΔ_T3 = (*ade6* SGA_T3 vs *ade6* SGA) − (Lib_T3 vs Lib). A ΔΔ value near zero means Torin1 exerts a proportionally equivalent effect on a given gene in both backgrounds: the gene responds the same way regardless of which background it is in. Departures from zero identify genes for which the drug effect is genuinely background-specific, pinpointing the subset of the genome where the *ade6* SGA control fails as a normalisation reference under the specific pharmacological condition applied.

Comparison of ΔΔ_T2 and ΔΔ_T3 (Figure 2D) showed only weak correspondence between doses (r = 0.35, RMSE = 0.262), indicating that the background-specific component of the Torin1 response is itself dose-dependent and not simply amplified between doses. A total of 1,090 of 2,581 scored genes (42.2%) differed by more than log_10_(1.5) in their ΔΔ values between the two concentrations. The regression slope of 0.56 further indicates that the magnitude of background-specific drug effects does not scale proportionally with Torin1 concentration, consistent with a non-linear physiological transition in which the *ade6* background becomes progressively and heterogeneously more sensitive to TOR inhibition as the dose increases. The broad, asymmetric ΔΔ_T3 distribution compared with ΔΔ_T2 is consistent with the emergence of a high-dose physiological state that the low-dose response does not predict. Together, the DiD analysis confirms and quantifies what the direct comparisons suggested: the *ade6* SGA background does not simply respond to Torin1 differently from the library; it responds in a fundamentally dose-rearranged manner that systematically compromises its use as a normalisation reference at pharmacologically active doses.

Comparing the dose-step responses directly (Lib_T3 vs Lib_T2 against *ade6* SGA_T3 vs *ade6* SGA_T2; Figure 2E), the two backgrounds showed very limited agreement (r = 0.22, RMSE = 0.224), with 892 of 2,458 genes (36.3%) exceeding the ±log_10_(1.5) threshold. The escalation from 2 μM to 3 μM Torin1 therefore produces fundamentally different gene-level fitness trajectories in the library and the *ade6* SGA background. The ranked ΔΔ_dose distribution (Figure 2F) showed that while most genes clustered near zero, a substantial subset showed strongly negative or positive values, indicating that the 2-to-3 μM escalation drives specific gene sets to more extreme background-specific responses. Together, these data demonstrate that the *ade6* SGA background diverges from the parental library in an increasingly non-linear, dose-dependent manner under Torin1 treatment, with control failure most evident at the higher drug concentration.

### Genes associated with control failure under Torin1 treatment

To characterise the genes driving divergence between the *ade6* SGA background and the haploid deletion library under Torin1 treatment, we generated heatmaps for genes selected from the positive and negative tails of the ΔΔ_T2, ΔΔ_T3, and ΔΔ_dose analyses (Figure 3A, 3B). These ΔΔ-based comparisons were used only for gene selection; the heatmaps display the corresponding interaction values of the selected genes across all nine contrast signatures.

**Figure 3.**
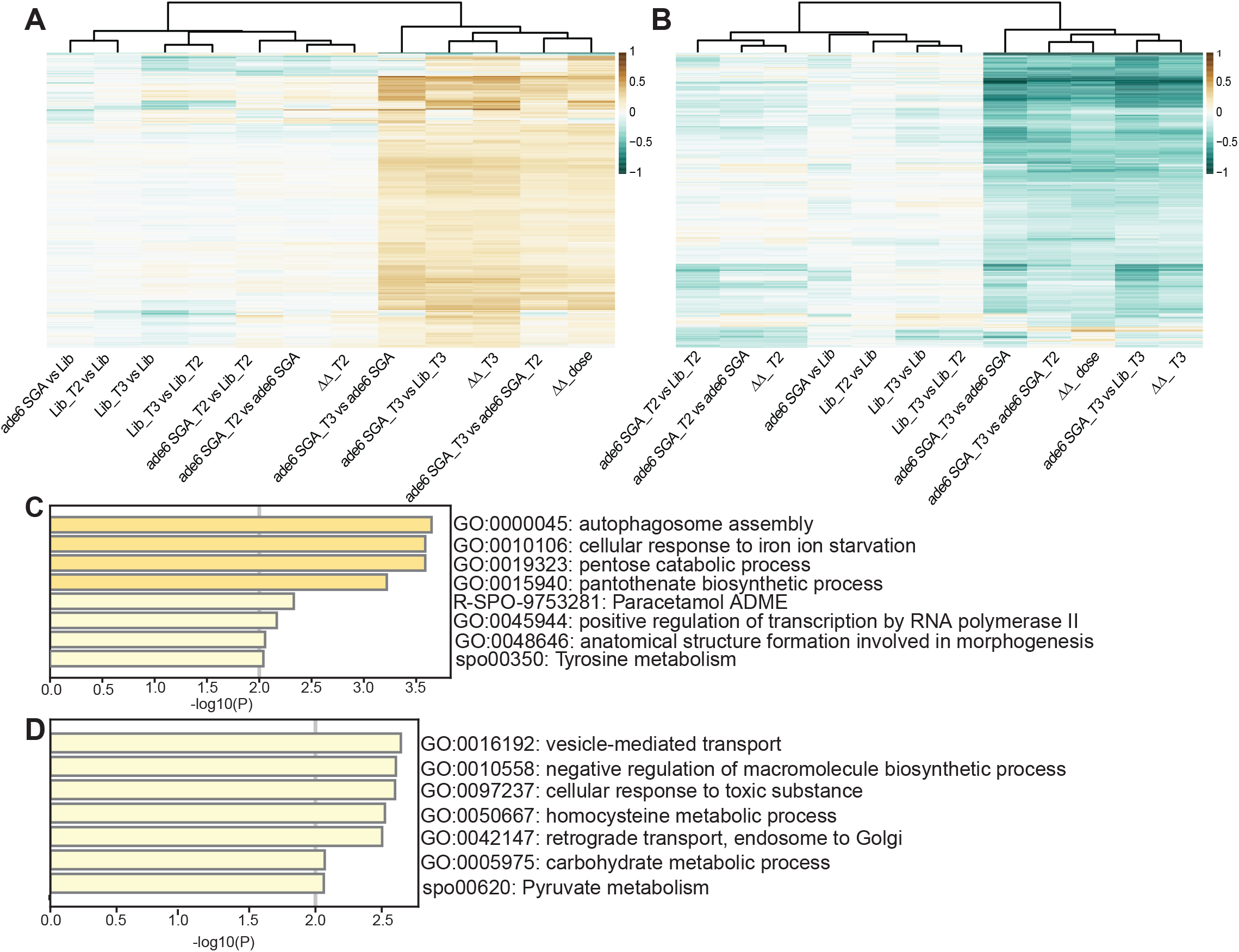
Heatmap and gene ontology enrichment analysis of genes associated with divergence between *ade6* SGA and the haploid deletion library under Torin1 treatment. (A) Heatmap of genes selected from the positive gene sets in Figure 2D (ΔΔ_T2 and ΔΔ_T3) and Figure 2F (ΔΔ_dose). (B) Heatmap of genes selected from the negative gene sets in Figure 2D (ΔΔ_T2 and ΔΔ_T3) and Figure 2F (ΔΔ_dose). Genes were selected using interaction thresholds of ≥ +0.176 for positive genes and ≤ -0.176 for negative genes. Rows represent genes and columns represent the baseline, Torin1-treatment, and derived contrast signatures. Values are shown on a centred colour scale, with negative values in teal, values near zero in white, and positive values in brown. (C) Gene ontology enrichment analysis of the positive gene set shown in panel A. (D) Gene ontology enrichment analysis of the negative gene set shown in panel B. Bars show enriched terms ranked by - log_10_ (P).

The positive-gene heatmap showed that the strongest positive interaction signals were concentrated almost exclusively in the T3-associated *ade6* contrasts -*ade6* SGA_T3 vs *ade6* SGA, *ade6* SGA_T3 vs Lib_T3, and *ade6* SGA_T3 vs *ade6* SGA_T2-while the untreated comparison and all library-centred contrasts remained close to zero across the same gene set. This pattern indicates that the positive divergence of these genes is specifically associated with the *ade6* background under 3 μM Torin1 treatment and is not a generalised response to the drug per se (which would also be reflected in the library-centred contrasts). The negative-gene heatmap showed the complementary pattern: the same T3-associated *ade6* contrasts dominated with the most strongly negative values, while the library-only contrasts and the untreated comparison remained comparatively muted. Together, both heatmaps confirm a clear dose-dependent, background-specific divergence that is concentrated in the 3 μM Torin1 condition and cannot be attributed to pre-existing baseline differences between backgrounds, further supporting the conclusion that high-dose TOR inhibition specifically uncovers functional non-equivalence of the *ade6* allelic states.

Gene ontology enrichment analysis revealed that the background divergence maps onto discrete, coherent biological programmes rather than a nonspecific genome-wide shift, indicating that the allelic difference between *ade6-M210/M216* and *ade6*Δ has pathway-specific rather than generalised consequences under TOR inhibition. Among the positively divergent genes -those more strongly induced or less inhibited in the *ade6* SGA background under high Torin1-the most enriched GO terms included autophagosome assembly, cellular response to iron ion starvation, pantothenate biosynthetic process, and pentose catabolic process (Figure 3C). These terms implicate autophagy induction, micronutrient sensing, and alternative carbon-source utilisation as processes differentially activated in the *ade6* SGA background, all of which are known to be regulated by TOR signalling and sensitive to the nutrient and metabolic context of the cell. Among the negatively divergent genes -those more strongly suppressed in the *ade6* SGA background under high Torin1-enriched terms included pyruvate metabolism, carbohydrate metabolic process, retrograde transport from endosome to Golgi, homocysteine metabolic process, and cellular response to toxic substance (Figure 3D). These terms point to suppression of central carbon metabolism and perturbation of intracellular vesicle trafficking specifically in the *ade6* SGA background under high-dose TOR inhibition. Collectively, the GO enrichment patterns are highly consistent with the known consequences of TOR inhibition in fission yeast, which rewires nutrient-responsive and stress-adaptive programmes in a metabolic-context-dependent manner^20,21^. The biological coherence and specificity of the enriched terms strongly support the interpretation that the *ade6-M210/M216* alleles in the library background confer a distinct metabolic context -one that renders the *ade6* SGA non-neutral for a substantial, biologically interpretable, and pharmacogenomically relevant subset of the genome under TOR inhibition.

## Discussion

Genetic interaction screens in yeast have transformed our understanding of gene function, cellular organisation, and the molecular logic of complex phenotypes^22,23^. The systematic mapping of synthetic lethal, suppressive, and quantitative interactions through SGA and related high-throughput methods has produced genome-scale interaction networks that reveal functional pathway architecture, define gene modules, and underpin predictive models of cellular function^2,5,24^. These networks have largely been built under standard laboratory growth conditions, where the genetic wiring of the cell is relatively stable and reproducible. The extension of SGA into pharmacogenomics -combining gene deletion with chemical or environmental perturbation-adds a powerful additional dimension: it can reveal which genetic states confer drug sensitivity or resistance, identify synthetic lethal opportunities for therapeutic exploitation, and illuminate mechanisms of drug action and off-target effects^25^. In fission yeast, pharmacogenomic SGA screens have been applied to TOR pathway inhibitors^12,13^, DNA-damaging agents^17^, and metabolic perturbations, revealing genetically encoded determinants of drug response that are often conserved across eukaryotes and relevant to human disease biology. A critical but underappreciated requirement for such screens is that the metabolic and genetic state of the normalisation control must remain appropriate under the specific pharmacological or stress condition applied -a requirement that the present study shows is not always met.

The central finding of this study is that the *ade6* background of a fission yeast deletion library -carrying the classical point mutations *ade6-M210* or *ade6-M216*^19,26^ rather than a complete deletion-is not biologically equivalent to the *ade6Δ* state of the SGA query strain, and this non-equivalence becomes functionally significant under TOR inhibition. The *ade6-M210/M216* alleles may confer adenine auxotrophy but retain the *ade6* locus with potentially residual enzymatic activity; fission yeast methods literature has long noted that these alleles are phenotypically distinct from null deletions and are “not as dark as a null allele”^27^. The *ade6* SGA background, by contrast, carries a deletion of *ade6*. These distinct allelic states create differences not only in adenine biosynthesis capacity but in the broader metabolic and regulatory landscape, differences that are cryptic under standard growth but unmasked by TOR inhibitors and possibly other drugs, compounds or stressors. Indeed, under nutrient-rich conditions these differences are apparently buffered, and the *ade6* SGA behaves similarly to the parental library (Figure 1E–F), functioning as a practical negative control for standard screens. The situation changes substantially when TOR signalling is inhibited. Torin1^28^ is an ATP-competitive inhibitor of both TORC1 and TORC2, inducing in fission yeast a nutrient-deprivation-like state associated growth suppression, metabolic stress, altered nutrient sensing, and activation of stress-adaptation pathways^29–31^. Under these conditions, previously ‘masked’ physiological differences between the *ade6-M210/M216* mutant alleles and a complete *ade6* deletion become functionally important, driving the dose-dependent divergence we describe here.

We propose that TOR inhibition places escalating pressure on nutrient adaptation, proteostasis, autophagy, and metabolic homeostasis, thereby exposing latent differences between the *ade6-M210/M216* and *ade6Δ* states. To minimise confounding by local chromosomal effects, genes within 100 kb of the *ade6* locus were excluded from all analyses^13^; the observed genome-wide divergence therefore reflects trans-acting metabolic and signalling consequences of the distinct allelic states. The dose-dependency is striking: at 2 μM Torin1 the two backgrounds retained moderate similarity (r = 0.56) with only 12.2% of genes exceeding the ±log_10_(1.5) threshold (Figure 2A), whereas 3 μM Torin1 collapsed this correspondence (r = 0.39) and drove 43.2% of genes beyond the same threshold (Figure 2B). The ΔΔ_T2 vs ΔΔ_T3 comparison (r = 0.35; Figure 2D) and the direct dose-escalation contrast (r = 0.22; Figure 2E) both reinforced this non-linearity, with the ranked ΔΔ_dose distribution developing broad background-specific tails at 3 μM Torin1 (Figure 2F). Together, these data indicate that increasing TOR inhibition drives cells across a physiological threshold at which background-specific effects become highly non-linear and pathway-dependent.

The GO enrichments further support a nutrient-homeostasis and stress-adaptation model. Positively divergent genes implicate autophagy (autophagosome assembly), iron starvation responses, and pantothenate biosynthesis -processes more active in the *ade6* SGA background under high Torin1. Negatively divergent genes point to suppressed vesicle-mediated transport, retrograde endosome-to-Golgi trafficking, and carbohydrate metabolism, consistent with TOR-dependent metabolic remodelling. Prior work showed that TOR inhibition in fission yeast globally rewires nutrient-responsive and stress-responsive programmes^20^, and that genetic interaction networks are extensively condition-dependent^17,32^, supporting the idea that the *ade6* allelic background is not neutral under pharmacological perturbation. Together, these findings indicate that while the *ade6* SGA functions as an appropriate control under standard growth conditions, it cannot be assumed neutral under TOR inhibition or other pharmacological stressors, and that condition-appropriate normalisation strategies should be adopted in pharmacogenomic SGA experiments.

Beyond the specific case of *ade6* and Torin1 described here, our findings illustrate a more general principle relevant to the design and interpretation of pharmacogenomic genetic interaction screens. Any control strain whose genetic background differs from the reference library -even at a locus considered metabolically neutral-carries an implicit assumption that the difference is functionally inert under the experimental condition applied. This assumption commonly holds under standard growth conditions but can fail when pharmacological or stress-induced perturbations alter the metabolic environment in ways that render previously silent allelic differences, functionally significant. Markers embedded in library backgrounds may be particularly vulnerable when screens are performed under conditions that perturb the pathways in which those markers participate. The fission yeast genetic interaction field, like that of budding yeast, relies on well-characterised marker strains whose behaviour has been validated, in the vast majority of cases, under benign growth conditions; the present study is a reminder that this validation does not automatically extend to pharmacogenomic conditions. More broadly, as SGA and related approaches are applied to an expanding range of compounds -including clinically relevant drugs, antimicrobial agents, nutrient-sensing pathway inhibitors, and environmental stressors-the genetic interaction landscape itself may be extensively rewired^32–34^, making the choice and validation of normalisation controls increasingly consequential for the reliability of the resulting maps.

Practically, we recommend two normalisation strategies for pharmacogenomic SGA screens. First, where feasible, using a drug- or stressor-treated library as the normalisation reference removes condition-specific background effects that may diverge between the library and the query control under treatment. Second, running a parallel untreated query SGA screen and using it as the reference for the drug-treated screen allows condition-specific genetic interactions to be isolated by subtraction, without requiring the entire library to be treated with the compound. Both approaches are substantially more reliable than using an untreated *ade6* SGA as the sole normalisation reference when the experimental condition includes pharmacological perturbation. We provide a quantitative framework, based on pairwise contrast correlation, difference-in-differences analysis, and GO enrichment of divergent genes that can be applied to assess normalisation control suitability in future screens. As chemical-genetic SGA approaches expand to new model systems, drug classes, and environmental conditions, empirical validation of normalisation controls under each specific experimental context will be essential for generating reproducible and biologically interpretable genetic interaction maps.

## Materials and Methods

### Media, reagents, and strains

Yeast extract with supplements (YES) medium, Edinburgh minimal medium (EMM), and agar were purchased from Formedium, UK. Adenine, uracil, and L-leucine were purchased from Merck/Sigma-Aldrich. Torin1 was purchased from MedChemExpress. Synthetic Genetic Array (SGA) analysis was performed using the Bioneer version 5.0 haploid deletion library constructed in the *ade6*-M210 or *ade6*-M216 ura4-D18 leu1-32 background^19^. The *ade6*Δ::natMX4/6 query strain was obtained from our laboratory strain collection.

### Growth conditions and SGA screening

The Bioneer deletion library^19^ and the *ade6Δ::natMX6* query strain were revived from -80 °C glycerol stocks and grown on YES medium. SGA manipulations were performed according to the quantitative SGA protocol described by Baryshnikova et al.^5^ High-density colony arraying and replica pinning were performed using a Rotor HDA robotic pinning system (Singer Instruments, Roadwater, UK). Drug screening was conducted in a 1536-density colony format, with each strain represented as quadruplicate colonies on each plate.

For Torin1 treatment experiments, library and *ade6* SGA plates were grown on YES medium in the presence or absence of Torin1 at the indicated concentrations. Colony images were acquired after incubation and colony-size measurements were obtained using the R package *gitter*^35^.

### Colony-size normalization and interaction analysis

All data are provided in Supplemental Data File 1. Quantitative interaction analysis between the haploid deletion library and the *ade6* SGA background under untreated and Torin1-treated conditions was performed using an in-house SGA analysis pipeline. Colonies smaller than 100 pixels in the corresponding control condition were excluded from downstream analysis. Strains missing because of technical or imaging errors were also removed.

Interaction values were calculated as:

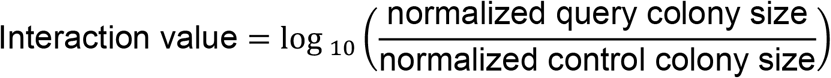

where normalized query colony size represents the colony size in the experimental condition and normalized control colony size represents the corresponding control condition. Interaction values ranged from +2 to −2.

To minimise potential linkage effects associated with the *ade6* locus, genes located within 100 kb of the *ade6* genomic region were excluded from the analysis in both the haploid deletion library and *ade6* SGA datasets.

For downstream comparative analyses and visualization, genes differing by more than ±0.176 interaction units were considered to exceed a 1.5-fold difference threshold, since log_10_(1.5) ≈ 0.176. Genes differing by more than ±0.301 interaction units exceeded a 2-fold difference threshold as log_10_(2) ≈ 0.301.

### Correlation, clustering, and principal component analysis

Pairwise similarity between contrast signatures was quantified using Pearson correlation coefficients:

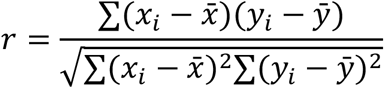

Hierarchical clustering was performed using correlation distance:

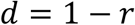

Where r is the Pearson correlation coefficient. Principal component analysis (PCA) was performed on gene-wise standardized interaction matrices.

### Identity scatter plots, RMSE and regression analysis

For identity scatter plots, the dashed black line represents the identity relationship:

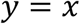

Deviation from the identity relationship was quantified using root mean square error (RMSE):

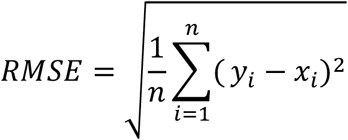

Lower RMSE values indicate closer agreement with the identity line, whereas higher RMSE values indicate greater deviation between contrasts.

Linear regression slopes were calculated using least-squares regression to assess whether the magnitude of response scaled equally between the two compared contrasts. A slope close to 1 indicates similar scaling, whereas slopes above or below 1 indicate systematic differences in response magnitude between contrasts.

### Difference-in-differences analyses

Difference-in-differences (ΔΔ) analyses were calculated as:

ΔΔ_T2 = (ade6 SGA_T2 vs ade SGA) - (Lib_T2 vs Lib)

ΔΔ_T3 = (ade6 SGA_T3 vs ade SGA) - (Lib_T3 vs Lib)

Dose-dependent divergence was calculated as:

ΔΔ_dose = (ade6 SGA_T3 vs ade6 SGA_T2) - (Lib_T3 vs Lib_T2)

### Gene ontology enrichment analysis

Gene ontology enrichment analyses were performed separately for positive and negative ΔΔ-associated gene sets. Enriched terms were ranked using -log_10_(P) values.

## Author Contributions

Rowshan Islam: Investigation, Validation, Formal analysis, Visualization, Writing

Olga Xintarakou: Investigation, Validation, Visualization

Charalampos Rallis: Conceptualization, Funding acquisition, Project administration, Resources, Supervision, Writing - original draft

